# Ribosomal RNA 2’-O-methylations regulate translation by impacting ribosome dynamics

**DOI:** 10.1101/2021.09.18.460910

**Authors:** Sohail Khoshnevis, R. Elizabeth Dreggors-Walker, Virginie Marchand, Yuri Motorin, Homa Ghalei

## Abstract

Protein synthesis by ribosomes is critically important for gene expression in all cells. The ribosomal RNAs (rRNAs) are marked by numerous chemical modifications. An abundant group of rRNA modifications, present in all domains of life, is 2’-O-methylation guided by box C/D small nucleolar RNAs (snoRNAs) which are part of small ribonucleoprotein complexes (snoRNPs). Although 2’-O-methylations are required for proper production of ribosomes, the mechanisms by which these modifications contribute to translation have remained elusive. Here, we show that a change in box C/D snoRNP biogenesis in actively growing yeast cells results in the production of hypo 2’-O-methylated ribosomes with distinct translational properties. Using RiboMeth-Seq for the quantitative analysis of 2’-O methylations, we identify site-specific perturbations of the rRNA 2’-O-methylation pattern and uncover sites that are not required for ribosome production under normal conditions. Characterization of the hypo 2’-O-methylated ribosomes reveals significant translational fidelity defects including frameshifting and near-cognate start codon selection. Using rRNA structural probing, we show that hypo 2’-O-methylation affects the inherent dynamics of the ribosomal subunits and impacts the binding of translation factor eIF1 thereby causing translational defects. Our data reveal an unforeseen spectrum of 2’-O-methylation heterogeneity in yeast rRNA and suggest a significant role for rRNA 2’-O-methylation in regulating cellular translation by controlling ribosome dynamics and ligand binding.

## Introduction

RNA molecules are subject to co- and post-transcriptional modifications which expand their chemical and topological properties (1, 2). Methylation of the 2’-O position of the ribose moiety of nucleotides is a highly abundant RNA modification, found in all four types of ribonucleotides in both coding and non-coding RNAs in all domains of life (3). Ribosomal RNAs (rRNAs) are a major target of ribose 2’-O methylations with fifty-five 2’-O-methylation sites identified in budding yeast and 106 in humans. Although rRNA 2’-O-methylations are critical for the proper production of ribosomes and accurate protein translation, their precise molecular contributions and mechanism of function are unknown (3–8). The chemical properties of 2’-O-methylations and the observations made based on their contributions to the structure of the ribosome, have suggested a role for rRNA modifications in local and global stabilization of the rRNA structure (1, 6, 7, 9, 10). Moreover, rRNA modifications contribute to the interactions of ligands with the ribosome (11, 12). Studies of modified nucleotides and RNA oligonucleotides have pointed to the importance of 2’-O-methylation in the stabilization of RNA structure by favoring the 3’-endo configuration of the ribose moiety and restricting the rotational freedom of the 3’-phosphate (13–18). However, how methylations of the rRNA backbone contribute to ribosome biogenesis and function remains largely unknown to date (4, 5, 19).

In eukaryotes and archaea, the box C/D small nucleolar RNAs (snoRNAs) guide the 2’-O-methylation of rRNA sites by base pairing to specific segments of the rRNA (20–25). Methylation is carried out by the action of the methyltransferase Nop1 (fibrillarin in humans) as part of a small nucleolar ribonucleoprotein complex (snoRNP). In this complex, four core proteins (Snu13, Nop56, Nop58, and Nop1) assemble on a snoRNA in a stepwise manner (26). This evolutionarily conserved assembly process requires the action of several assembly factors that confer tight regulation of the biogenesis and turnover of box C/D snoRNAs.

Recent quantitative studies have shown that 2’-O-methylations are not ubiquitously present on all of the cell’ s ribosomes, suggesting that these modifications may be an adjustable feature to fine tune the function of ribosomes (4, 27–35). However, because most rRNA 2’-O-methylations are deposited at an early stage during ribosome biogenesis, it is not clear how these modifications can be adjusted or removed and the source of rRNA 2’-O-methylation heterogeneity remains unknown (4, 5, 36). Attempts to investigate the repertoire of sub-stoichiometric 2’-O-methylation of rRNA include the study of natural variations between cell lines (31, 35), changes of rRNA 2’-O-methylations in primary human breast tumors (37), differential rRNA modifications during mouse development (38), changes of modifications in the presence or absence of the antitumor protein p53 (28, 34) which regulates the methyltransferase fibrillarin (39), changes caused by depletion of fibrillarin (27) and the use of a catalytically inactive Nop1 mutant (40). Surprisingly, however, the methods used to study the effect of rRNA hypo 2’-O-methylation have not allowed identification of all potential variable modification sites. For example, mutating the active site of Nop1, which is expected to directly affect rRNA 2’-O-methylations, did not change the methylation pattern of the ribosome (40). The alternative approach of knocking down fibrillarin using siRNA in human cell lines (27), allowed researchers to identify a set of 2’-O-methylations which are specifically reduced in the absence of fibrillarin. However, because cells depleted of fibrillarin hardly divide or make any new ribosomes and likely rely on their already assembled ribosomes for survival (41–44), limited modification changes were observed using this strategy. Furthermore, which effects of fibrillarin knockdown are direct versus indirect was not discerned (45). Therefore, obtaining a comprehensive view of the rRNA sites that have the potential to be missed, removed or regulated has not been possible thus far.

To overcome this limitation, we present here a new strategy which involves decreasing the biogenesis of box C/D snoRNAs via a single mutation in an essential box C/D snoRNP assembly factor, Bcd1. Bcd1 is an evolutionarily conserved protein that cooperates with several other assembly factors to direct the assembly of box C/D snoRNAs into functional snoRNP complexes (26). Bcd1 controls the steady-state levels of box C/D snoRNAs by regulating the assembly of the Nop58 protein into pre-snoRNPs and by mediating the interaction of Snu13 with snoRNAs (46–48). Using a variant of Bcd1 (*bcd1-D72A*), which causes a global box C/D snoRNA downregulation but allows cell growth and ribosome biogenesis (48), here, we reveal a wide spectrum of variable rRNA 2’-O-methylation sites in yeast ribosomes and identify their impact on translation. Strikingly, our data reveal that more than 70% of yeast rRNA 2’-O-methylation sites have the potential to be significantly hypomethylated. We also show that 2’-O-methylation affects the dynamics of the rRNA resulting in a change in the balance between different conformational states of the ribosomes required for translation. Finally, our data show that rRNA hypomethylation also impacts the binding of initiation factor 1 (eIF1) to the small ribosomal subunit. Together, these results allow to dissect those rRNA 2’-O-methylation sites that are required for ribosome biogenesis from those that are dispensable and may have other functional roles, and link rRNA 2’-O-methylation to specific features of the ribosomes.

## Results

### Heterogeneity of rRNA 2’-O-methylation sites in yeast rRNA reveals sites that are dispensable for ribosome biogenesis

To address which of the rRNA 2’-O-methylations sites are tunable in yeast and identify those that are dispensable for ribosome production in viable cells, we exploited a mutation in the modification machinery that alters the biogenesis of box C/D snoRNAs in actively growing yeast cells. For this purpose, we engineered the *bcd1-D72A* mutation in yeast cells using CRISPR/Cas9 genome editing (48). The introduction of this mutation into the genome ensures that all the ribosomes will be made in the presence of defective Bcd1, which causes cells to have low steady-state levels of box C/D snoRNAs (48). We then performed RiboMeth-Seq analysis (31) on RNAs isolated from actively growing wild-type control yeast cells and those expressing *bcd1*-D72A. Table S1 lists the average MethScores (ScoreC) for all rRNA methylation sites in wild-type control and mutant *bcd1-D72A* cells. In line with the previous observations (30, 31), 47 of the 54 2’-O-methylation sites in wild-type control yeast cells are methylated at high levels (with MethScore > 0.8) and only a small fraction (7/54 sites) show MethScores below 0.8. While the average MethScore for wild-type control is 0.85, analysis of data from *bcd1-D72A* cells reveal a much lower level of methylation (average MethScore 0.43). These data indicate that the majority of rRNA 2’-O-methylation positions are hypomethylated in *bcd1-D72A* cells.

Analysis of the 2’-O-methylation sites in the 18S rRNA of *bcd1-D72A* cells reveal 7 stable sites with similar methylation levels to the wild-type control (MethScore > 0.8), 8 variable sites (MethScore between 0.4 and 0.8), and 3 highly hypomethylated sites (MethScore < 0.4). The two most stable modification sites are found close to the decoding center (G1428 and C1639 with MethScore > 0.9) (Figure 1A). In contrast, the hypomethylated sites within 18S rRNA do not cluster at a specific region within. For 25S rRNA, we identify 6 stable sites with near-complete methylation, 15 variable sites, and 15 hypomethylated sites. Interestingly, although the peptidyl transferase center (PTC) and the tRNA accommodation corridor are devoid of 2’-O-methylated sites, several hypomethylated or variable sites neighbor these regions (Figure 1B). Taken together, our RiboMeth-Seq data reveal that while 24% (13/54) of rRNA 2’-O-methylation sites are essential for ribosome biogenesis and function, while other rRNA 2’-O-methylations can be variable (42%, 23/54) or even dispensable (33%, 18/54).

**Figure 1.**
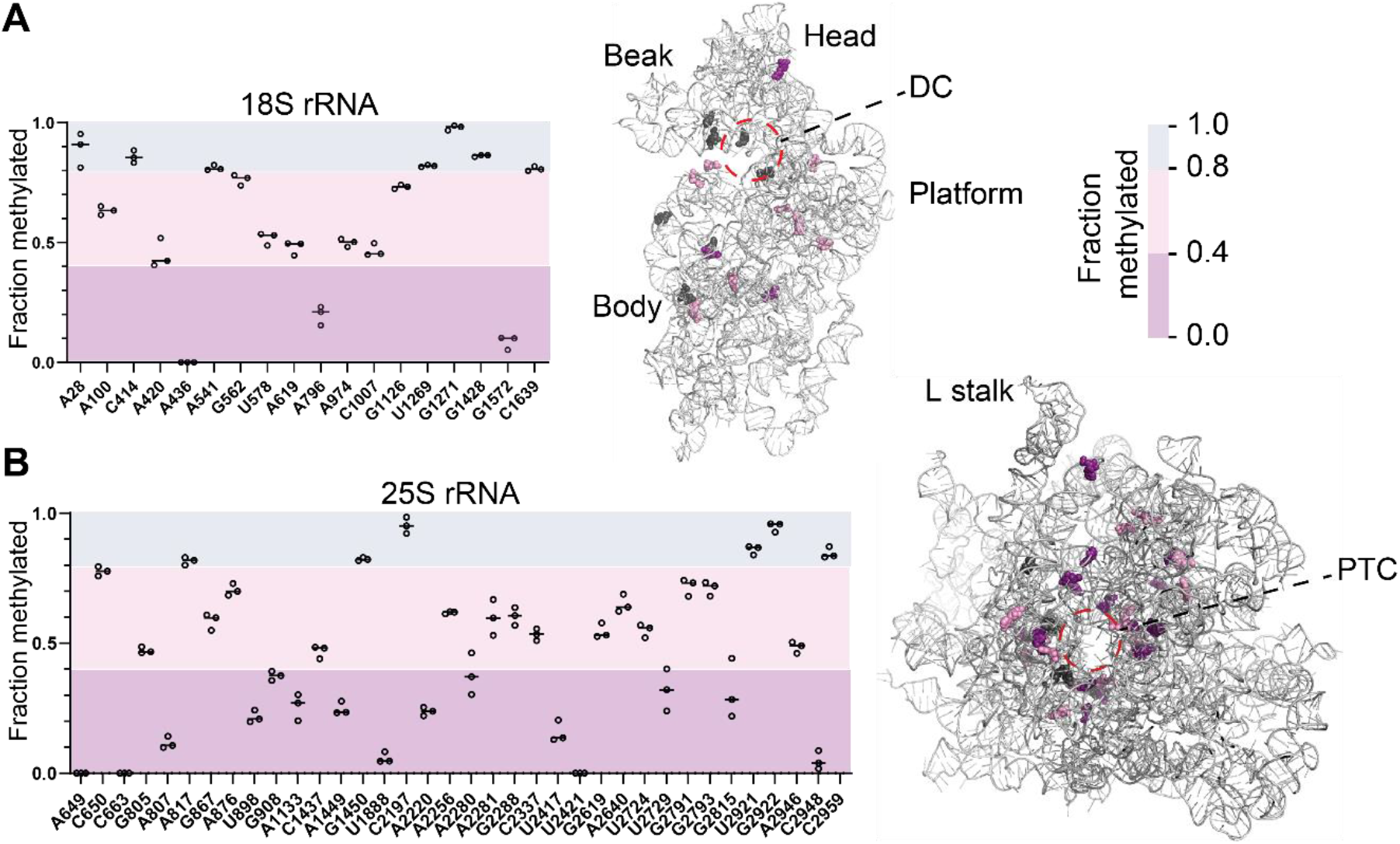
rRNA 2’-O-methylation sites change in a site-specific manner in *bcd1-D72A* cells. The fraction of 2’-O-methylation (MethScore) at each modification site in 18S (**A**) and 25S (**B**) rRNA in *bcd1-D72A* relative to wild-type control cells was evaluated by RiboMeth-Seq. Data are expressed as mean MethScore values ± SD (n=3 independent biological replicates). The position of each modification is marked on the 18S (**A**) and 25S (**B**) rRNA structure (PDB ID: 6GQV). The stable sites (MethScore >0.8) are colored in grey, the variable sites (0.4 < MethScore < 0.8) in pink, and the hypo 2’-O-methylated sites (MethScore < 0.4) in magenta. DC stands for decoding center and PTC for peptidyl transferase center.

### rRNA hypomethylation affects the fidelity of protein synthesis

Ribosomal RNA modifications are critically important for the function and fidelity of ribosomes (7, 11, 49, 50). We, therefore, tested whether the change of rRNA 2’-O-methylation pattern in *bcd1-D72A* cells affects the efficiency and accuracy of protein synthesis. To assess the translational efficiency of hypo 2’-O-methylated ribosomes, we analyzed the incorporation rate of L-homopropargylglycine (HPG), an amino acid analog of methionine containing an alkyne moiety that can be fluorescently modified, into newly synthesized peptides in rapidly dividing wild-type or *bcd1-D72A* cells. HPG-containing proteins were fluorescently labeled by the addition of Alexa Fluor 488 and separated from the unincorporated dye by SDS-PAGE (Figure 2A). Quantification of the levels of newly synthesized proteins relative to the total proteins in wild-type and *bcd1-D72A* cells reveal that while the total protein synthesis is higher in wild-type control cells, the rate of HPG incorporation over time is not significantly different between the two strains (Figure 2B). However, we observe a higher fluorescent signal at each time point in control cells versus *bcd1-D72A* cells, suggesting a higher number of translating ribosomes in wild-type control cells in agreement with the ribosome biogenesis defects we had previously observed in *bcd1-D72A* cells (48).

**Figure 2.**
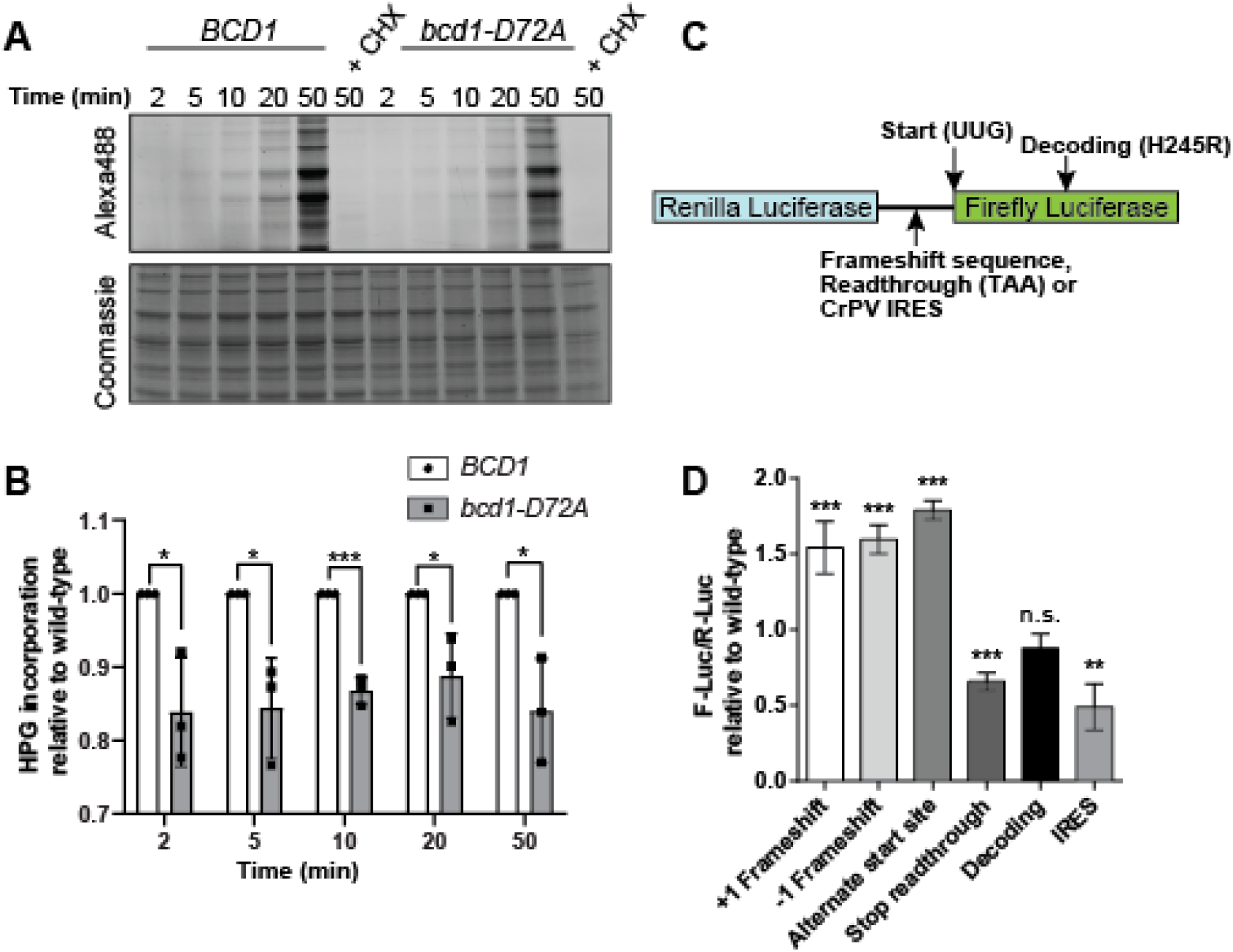
rRNA hypomethylation affects the function and fidelity of ribosomes. (**A**) Analysis of the incorporation rate of L-homopropargylglycine (HPG) into newly synthesized peptides in rapidly dividing yeast cells expressing wild-type or mutant Bcd1 over a time course (2-50 min). HPG-containing proteins were fluorescently labeled by addition of Alexa Fluor 488 and separated from unincorporated dye by SDS-PAGE and imaged (top) before staining with Coomassie blue for total protein detection (bottom). (**B**) Quantification of the data shown in panel A. At each time point, the ratio of the newly synthesized protein to the total protein for *bcd1-D72A* is normalized to the wild type. (**C**) Schematic of the dual luciferase plasmids used in this study. For all plasmids, Renilla luciferase is constitutively expressed, while the expression of firefly luciferase is dependent on a specific translational defect/element. (**D**) Expression of firefly and Renilla luciferase was measured in wild-type control or *bcd1-D72A* yeast harboring the indicated plasmids. The ratio of firefly luciferase to Renilla luciferase was normalized to the control plasmids and shown relative to wild-type. Significance for each column was analyzed relative to the wild-type control with multiple t-test comparisons. n.s., non-significant; *p ≤ 0.05; **p ≤ 0.01; ***p ≤ 0.001.

Because changes in ribosome number and/or composition can affect the accuracy of protein synthesis and impact the ability of ribosomes to initiate from internal ribosome entry sites (IRESs) (27, 51–53), we next assayed the fidelity of translation in wild-type control and *bcd1-D72A* cells. For this purpose, we used a set of established dual-luciferase reporter plasmids in which the translation of the firefly luciferase is affected by −1 and +1 programmed frameshifting, alternate start codon selection, stop codon readthrough, miscoding or initiation from an IRES element (Figure 2C, (54–58)). As an internal control, all plasmids contain a constitutively expressed Renilla luciferase used for normalization. This analysis reveals that, compared to control ribosomes, ribosomes from *bcd1-D72A* cells are significantly defective in near-cognate start codon selection and frameshifting (Figure 2D). The data also reveal that rRNA hypo 2’-O-methylation impacts the intrinsic capability of ribosomes to initiate translation from IRES elements and recognition of the stop codon.

### rRNA hypo 2’-O-methylation impacts the rotational status of ribosomes

Recent work has shown that the mRNA structures that promote programmed frameshifting in bacteria change the reading frame of the ribosome by increasing the rotated-state pause (59–61), thus providing a link between ribosome dynamics and frameshifting. Because 2’-O-methylation affects the flexibility of RNA, we hypothesized that the observed changes in mRNA frameshifting in *bcd1-D72A* cells could arise from altered dynamics of the ribosome. To address this question, we used RNA structure probing. Several key residues in both the small and large ribosomal subunits undergo detectable changes in conformation upon transition between rotated and non-rotated ribosomal states (62). The most prominent of these sites is G913 in 18S rRNA (SSU-G913) which is located at the inter-subunit bridge B7a. To compare the rotational dynamics of the wild-type control and hypomethylated ribosomes, we probed this nucleotide in wild-type and *bcd1-D72A* cells using phenylglyoxal (PGO) (Figure 3A). As a reference, we took advantage of two mutations in *RPL3* (*rpl3-W255C* and *rpl3-H256A*), that are known to stabilize ribosomes in non-rotated and rotated states, respectively (63). We then compared the rotational status of wild-type and hypo 2’-O-methylated ribosomes to the rotated and non-rotated ribosomes from *RPL3* mutant cells (Figure 3B).

**Figure 3.**
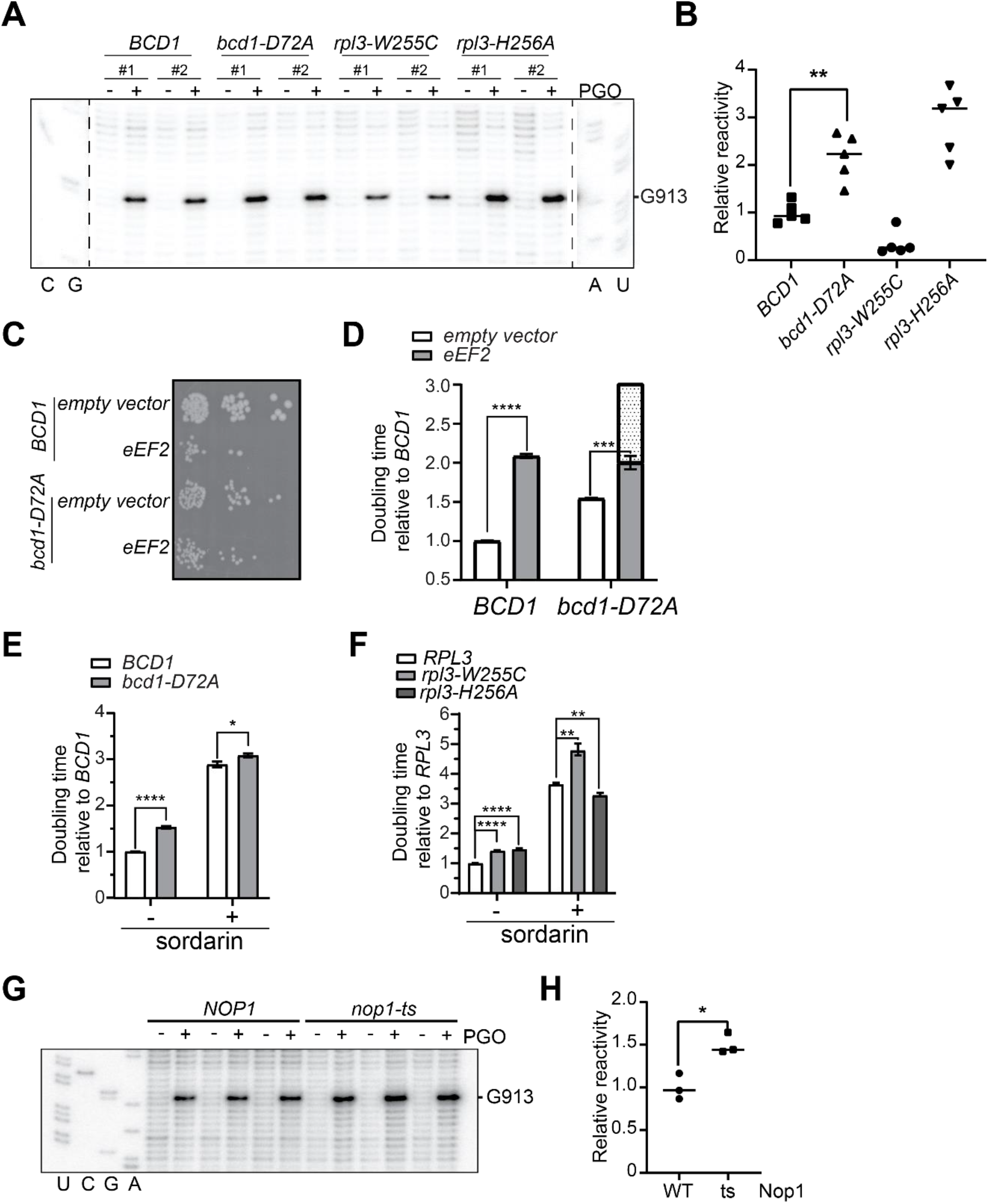
Hypo 2’-O-methylated ribosomes are found more in the rotated conformation. (**A**) *In vivo* RNA structure probing of cells expressing either wild-type or D72A variant of Bcd1 with or without PGO treatment to probe the accessibility of SSU G913. Cells expressing W255C or H256A variants of Rpl3 were used as control for rotation status. Two biological replicates are shown in the figure. (**B**) Quantification of the SSU G913 modification by PGO. (**C**) *bcd1-D72A* cells are less sensitive to the overexpression of eEF2 than wild type cells. **(D)** quantification of the growth of BCD1 and bcd1-D72A expressing eEF2. Dot pattern indicates the expected doubling time of *bcd1-D72A* if there was no rescue of the *bcd1-D72A* by eEF2 overexpression. (**E**) *bcd1-D72A* cells are less sensitive to sordarin (3 μg/mL) than wild type cells. (**F**) Whereas *rpl3-W255C* has the same sensitivity to sordarin as the wildtype, *rpl3-H256A* shows less sensitivity to sordarin. (**G**) Probing the accessibility of SSU G913 in *NOP1* and *nop1-ts* using PGO. Three biological replicates are shown in the figure. (**H**) Quantification of the SSU G913 modification by PGO. Significance was analyzed relative to the wild-type control with multiple t-test comparisons. *p ≤ 0.05; **p ≤ 0.01; ***p ≤ 0.001; ****p≤ 0.0001.

Our analyses reveal that *in vivo*, the SSU-G913 nucleotide in *rpl3-W255C* ribosomes is protected and show minor reactivity to PGO. In *rpl3-H256A* cells, however, in which the ribosomes are stabilized in the rotated state, the SSU-G913 nucleotide is more accessible when probed with PGO. Comparing the reactivity of SSU-G913 in wild-type control and *bcd1-D72A* cells with those of *rpl3-W255C* and *rpl3-H256A* reveal that the hypomethylated ribosomes from *bcd1-D72A* cells are at least as accessible to PGO as the rotated ribosomes from *rpl3-H256A cells* suggesting that the majority of ribosomes from *bcd1-D72A* cells are in the rotated conformation relative to the wild-type control (Figure 3A, B). These data suggest that ribosomes from *bcd1-D72A* cells favor the rotated state and that the increased rate of frameshifting in hypomethylated ribosomes could be due to the increased time the hypo 2’-O-methylated ribosomes spend in the rotated state.

Elongation factor 2 (EF-G in bacteria and eEF2 in eukaryotes) binds to pre-translocation ribosomes and stabilizes the rotated state (64). Mutations in yeast ribosomes which stabilize the rotated conformation increase the affinity of eEF2 for ribosomes (62, 63). Because the overexpression of eEF2 impairs yeast growth (Figure 3C, D), we hypothesized that as the hypo 2’-O-methylated ribosomes preferentially assume the rotated conformation in the cell, the overexpression of eEF2 should have less deleterious effect on *bcd1-D72A* cells compared to the wild-type control. To test this, we compared the growth of wild-type control and *bcd1-D72A* yeast cells overexpressing eEF2. While overexpression of eEF2 causes a severe growth defect in wild-type cells, it does not impact the growth of *bcd1-D72A* cells to the same extent (Figure 3C). To further quantify this growth difference, we measured the growth of wild-type control and *bcd1-D72A* yeast cells overexpressing eEF2 cells during their logarithmic phase of growth. These data indicate that indeed *bcd1-D72A* cells are less sensitive to the overexpression of eEF2 than the control cells (Figure 3D).

To further test whether hypomethylated ribosomes spend more time in the rotated state, we measured the growth of *bcd1-D72A* and control cells in the presence of sordarin. Sordarin is an inhibitor of the eEF2 that binds to ribosome-bound eEF2 and prevents its domain movements which are required for translocation (65). We analyzed the effect of sordarin on the growth of *bcd1-D72A* and wild-type control cells and compared that to the growth of reference cells expressing wild-type *RPL3*, *rpl3-W255C,* or *rpl3-H256A*. Whereas cells expressing wildtype *RPL3* and *rpl3-W255C* show similar sensitivity to sordarin, *rpl3-H256A* and *bcd1-D72A* cells are both less sensitive to sordarin (Figure 3E, F). These data further support the notion that hypo 2’-O-methylated ribosomes, similar to *rpl3-H256A* harboring ribosomes, spend more time in the rotated state. Taken together, these data provide the first evidence that the 2’-O methylation status of rRNA affects the ribosomal rotation state. Importantly, because there are no 2’-O-methylation sites near the SSU-G913 nucleotide, our data suggest that the ribosome rotational changes observed in *bcd1-D72A* cells are due to long-range effects.

Recent studies have proposed a function for snoRNAs in chaperoning the folding of rRNA (66). The levels of box C/D snoRNAs are significantly lower in *bcd1-D72A* cells relative to wild-type control cells (46, 48). The changes we detect in the conformation of ribosomes could therefore arise either directly from the lack of 2’-O-methylations or indirectly from the decreased levels of snoRNAs. To dissect the role of snoRNA binding and chaperoning from the effect of rRNA 2’-O-methylations on the ribosome structure, we took advantage of a temperature-sensitive (ts) mutation in the methyltransferase Nop1 (67). Cells harboring this mutation grow more slowly than wild-type control cells at 37°C, and have lower rRNA methylation levels but do not show any change in snoRNA levels (Figure S1). *In vivo* analysis of the SSU-G913 modification by PGO in *nop1-ts* cells compared to wild-type control cells show higher accessibility of SSU-G913 in *nop1-ts* cells, suggesting the stabilization of the rotated ribosomal conformation in the hypo 2’-O-methylated ribosomes (Figure 3G, H). Because the level of snoRNAs is similar to the wild-type control cells in the *nop1-ts* cells, these data indicate that the observed defects in the inherent dynamics of the ribosome in *bcd1-D72A* cells are more due to the decreased rRNA 2’-O-methylation rather than a decrease in snoRNA levels.

### Binding of eIF1 to the hypo 2’-O-methylated small ribosomal subunit is weakened *in vivo* and *in vitro*

Eukaryotic initiation factor 1 (eIF1) plays an important role in ensuring the selection of cognate start codon and antagonizing the near-cognate start codon selection by stabilizing the open, scanning-competent conformation of the small ribosomal subunit (40S) (68–70). Upon proper start codon selection, eIF1 is released from 40S allowing rearrangement of the ribosome from the open to closed conformation (55, 71). Because ribosomes from *bcd1-D72A* cells show an elevated level of near-cognate start codon recognition (Figure 2D), we hypothesized that the hypo 2’-O-methylated ribosomes from *bcd1-D72A* cells cannot bind to eIF1 as efficiently as wild-type ribosomes. To test this hypothesis, cells expressing wild-type or the D72A variant of Bcd1 were fixed with formaldehyde in the mid-log phase and ribosome-bound and free eIF1 were separated by centrifugation on sucrose density gradients and quantified by western blotting. While ~25% of the eIF1 co-migrate with the ribosomes in wild-type control cells, the binding of eIF1 to the hypo 2’-O-methylated ribosomes in *bcd1-D72A* cells is significantly reduced (Figure 4A).

**Figure 4.**
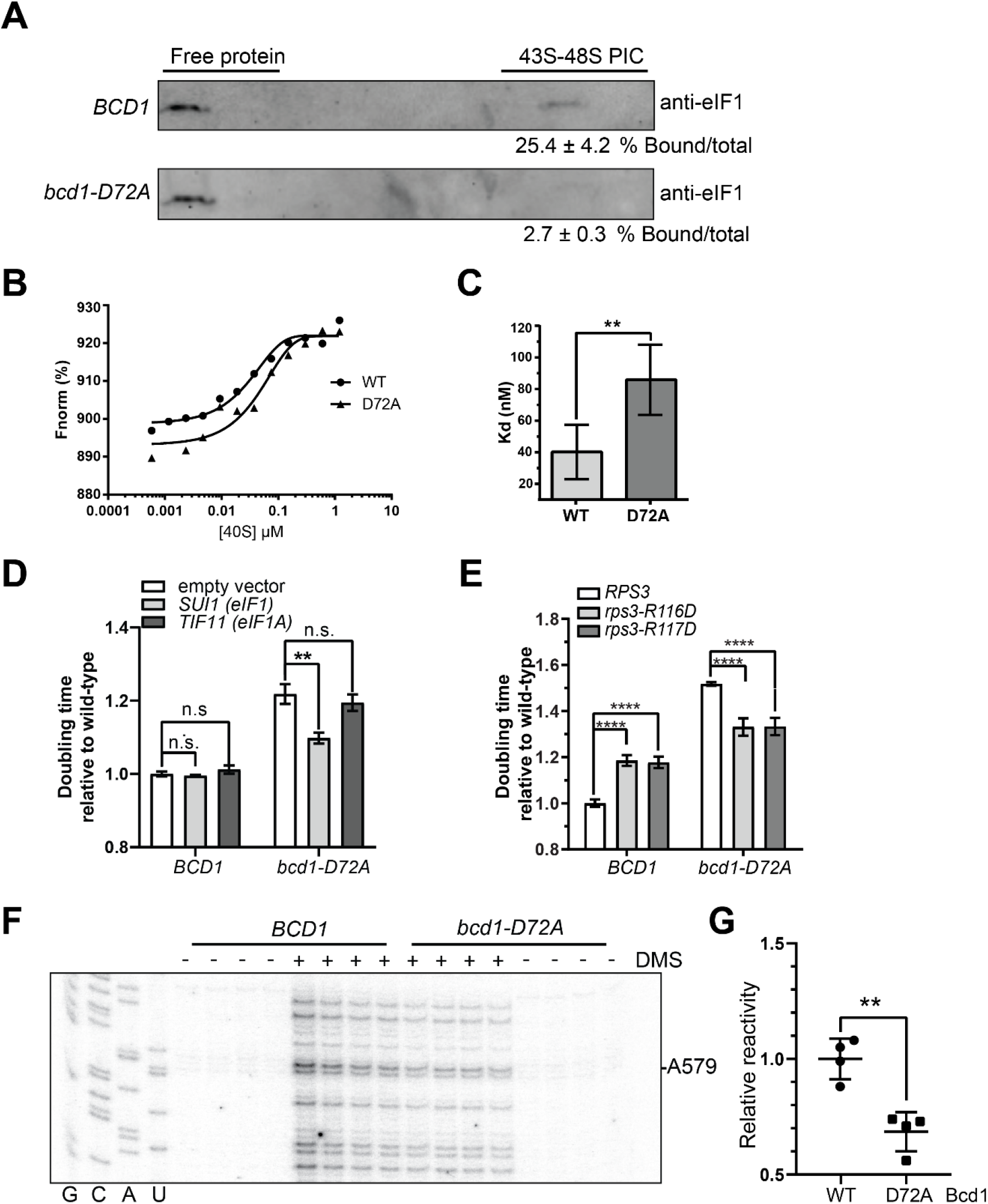
Binding of eIF1 to hypomethylated 40S is weakened *in vivo* and *in vitro*. (**A**) Western blot against eIF1 from sucrose gradients of formaldehyde-fixed whole-cell extracts from *BCD1-WT* (top) or *bcd1-D72A* (bottom). The ratio of eIF1 in 43-48S PIC relative to the total eIF1 is depicted under each blot. (**B**) Changes in fluorescence plotted against the 40S concentration. Data were fitted with a non-linear regression model in GraphPad Prism 8.0. (**C**) Dissociation constants calculated from 4 independent experiments as shown in panel B. (**D**) Comparison of doubling times of *BCD1-WT* and *bcd1-D72A* harboring either an empty vector or vectors expressing *SUI1* (eIF1) or *TIF11* (eIF1A) in minimal medium containing glucose. The data are the average of four biological replicates, and error bars show SEM. Each column was analyzed relative to the *BCD1-WT* with empty vector. (**E**) Comparison of doubling times of *BCD1-WT* and *bcd1-D72A* in which the endogenous *RPS3* gene is deleted and either WT, R116D or R117D variant of *RPS3* are expressed from a plasmid. The data are the average of four biological replicates, and error bars show SEM. Each column was analyzed relative to the *BCD1-WT* with *RPS3-WT* plasmid. (**F**) Probing the accessibility of SSU A579 in 40S ribosomal subunits purified from *BCD1* and *bcd1-D72A* using DMS. Four biological replicates are shown in the figure. (**G**) Quantification of the SSU A579 modification by DMS. Significance was analyzed relative to the wild-type control with multiple t-test comparisons. n.s., non-significant; **p ≤ 0.01; ***p ≤ 0.001; ****p≤ 0.0001.

To corroborate this finding, we next assessed the binding of eIF1 to 40S ribosomal subunits isolated from wild-type control and *bcd1-D72A* cells. For this purpose, we fluorescently labeled eIF1 and determined its binding affinity for purified 40S subunits. The results from these binding assays reveal that eIF1 binds to 40S subunits isolated from wild-type cells with a dissociation constant (Kd) of ~40 nM, comparable to previous reports (72). However, the affinity of eIF1 for hypo 2’-O-methylated 40S ribosomal subunits is lower by half to ~80 nM (Figure 4B-C). In line with this decrease in affinity, overexpression of eIF1, but not eIF1A, rescues the growth defects in *bcd1-D72A* cells to a great extent while it does not affect the wild-type control cells (Figure 4D).

The biochemical and genetic data above suggest that rRNA 2’-O-methylations play a key role in regulating the binding of eIF1 thereby ensuring the stringency of start codon selection. A prediction from this finding is that other factors/mutants that antagonize the near-cognate start codon selection could at least partially rescue the slow growth phenotype of *bcd1-D72A* cells. To test this idea, we took advantage of two *rps3* missense mutants (R116D and R117D) that destabilize the closed conformation of the 48S pre-initiation complex (PIC) and antagonize the near-cognate start codon selection (73). First, we replaced the endogenous promoter of the *RPS3* gene with a galactose-inducible/glucose repressible promoter in wild-type control and *bcd1-D72A* cells. This allowed us to compare the impact of expression of wild-type and mutant versions of *RPS3* on the growth of wild-type control and *bcd1-D72A* cells by transforming in plasmids encoding these variants and turning off the expression of endogenous *RPS3*. Interestingly, expression of both variants of *RPS3* rescued the slow growth defect of *bcd1-D72A* cells (Figure 4E).

The 18S rRNA folds into distinct domains known as head, shoulder, and body. The mRNA binding site lies within a cleft between the head and body domains. In the open, scanning competent conformation, the head and body domains are farther apart from each other compared to the closed conformation, allowing the rapid movement of the 40S on the mRNA. In the closed conformation, the head and body come close to each other, locking the initiation codon in the P-site (74, 75). A distinct structural feature of the closed state is the appearance of the mRNA latch composed of elements in h18 in the body with h34 and Rps3 in the head domain (69). Comparing the latch nucleotides in the closed and open conformations (76) suggests that the A579 nucleotide is more solvent accessible in the open conformation and hence more likely to be modified by RNA structure probing reagents (Figure S2). To probe if the equilibrium between the open and closed conformations of 40S is affected in hypo 2’-O-methylated ribosomes, we treated 40S subunits isolated from control or *bcd1-D72A* cells to dimethyl sulfate (DMS) and analyzed the accessibility of the A579 nucleotide (Figure 4F). Our analysis shows a decreased reactivity of the A579 nucleotide towards DMS in *bcd1-D72A* cells relative to the control cells, suggesting that the mRNA latch is closed in a higher population of 40S ribosomes in *bcd1-D72A* relative to the wild-type control cells (Figure 4G).

Altogether, these data strongly suggest that rRNA hypo 2’-O-methylation changes the inherent conformational dynamics of ribosomes thereby impacting ribosome-factor interactions leading to translational errors. Thus, the deficiency in start codon selection in hypo 2’-O-methylated ribosomes may be attributed to the weaker binding of the ribosome to eIF1 and changes in ribosome dynamics.

## Discussion

snoRNA-guided ribose 2’-O methylation is an evolutionarily conserved common form of methylation in rRNA. Even though RNA 2’-O-methylation changes have been linked to a large number of human diseases (5, 8, 37, 39, 53, 77, 78), the role of rRNA 2’-O-methylations for the function of ribosomes is not understood to date. Recent findings indicate the plasticity of 2’-O-methylation in rRNA, proposing new avenues for fine-tuning the function of the ribosome (5, 27, 28, 35, 37, 38). Here, we describe a new strategy for surveying the repertoire of sub-stoichiometric rRNA 2’-O-methylation sites by exploiting a defect in box C/D snoRNP assembly. This approach allowed us to lower the overall level of box C/D snoRNAs thereby globally changing rRNA 2’-O methylations without severely affecting cell survival, enabling us to probe rRNA 2’-O-methylation changes that had remained elusive so far.

Interestingly, as suggested from previous studies of human snoRNAs (28), comparing the snoRNA levels to the fraction of modification of their corresponding 2’-O-methylation sites indicated that there was no direct correlation between the steady-state box C/D snoRNA levels and the methylation status of nucleotides. While the majority of box C/D snoRNA levels decreased by more than 60% in *bcd1-D72A* cells, in some cases we observed near complete 2’-O-methylation (Table S2). The observation that most stable methylation sites had their corresponding snoRNA levels reduced by 80-90%, suggests a threshold model for rRNA modification where the presence of even a small amount of snoRNAs is sufficient for 2’-O-methylation of the majority of transcribed rRNA.

Of the 54 2’-O-methylation sites in yeast rRNA, 38 are conserved between yeast and humans. The potential of these conserved sites for variability in their methylation was assessed previously after knockdown of the methyltransferase fibrillarin (27). This study showed that 6/38 of the conserved sites between yeast and humans were less than 80% methylated and could be substantially altered. With our approach of manipulating snoRNA levels in yeast, we identified 25/38 conserved sites in yeast that were methylated in less than 80% of the rRNA population (Figure S3). Interestingly, all but one of the 6 variable conserved sites that were identified in human rRNA had their yeast equivalents also hypomodified in the *bcd1-D72A* mutants. The Cm2197 site in yeast rRNA is the only exception which remains fully methylated in *bcd1-D72A* cells, yet its human equivalent shows significant hypomethylation in fibrillarin depleted HeLa cells (27). We attribute this difference to the resistance of the guiding snoRNA, snR76, to *bcd1-D72A* mutation (Table S2) or the higher stability of this snoRNA. A possible explanation for the higher number of variable modifications in yeast rRNA compared to human rRNA could result from organismal differences. However, as opposed to cells depleted of fibrillarin, which undergo limited division and do not make new ribosomes (43), *bcd1-D72A* yeast cells we have used in this study keep dividing and making new ribosomes (48). This can result in a higher rate of hypomodification and allow us to map sub-stoichiometric 2’-O-methylations in a more thorough way than was possible previously.

While most of the box C/D snoRNAs in yeast guide the site-specific modification of rRNA, a few snoRNAs play a role in rRNA folding and processing (79). Recently, the box H/ACA snR35 was proposed to prevent the premature folding of helix31 in pre-40S, thereby contributing to rRNA folding in a manner distinct from its modifying role (66). Given the decrease in snoRNA levels in *bcd1-D72A* cells relative to wild-type control, we tested whether the changes in the dynamics of the ribosomes arise from the lack of methylation or decreased snoRNA levels. Our results show that altered inherent ribosomal dynamics can be caused by changes in rRNA 2’-O-methylation but not by the loss of snoRNAs and their chaperoning effect. These findings corroborate previous work on several box C/D and H/ACA snoRNAs which point to the importance of rRNA modifications in addition to the chaperoning role of snoRNAs (50). However, further studies are required to dissect the potential role of snoRNAs in chaperoning rRNA folding and investigating the contributions from these events to rRNA dynamics.

Whereas deletion of most individual box C/D snoRNAs does not have a major effect on yeast cell growth under normal conditions (80–82), simultaneous deletion of groups of rRNA 2’-O-methylations or using a catalytically inactive Nop1 severely affects cell growth and results in translation defects (4, 42, 83). For example, the absence of modifications in 25S H69 as well as around the decoding center and the A-site finger causes translational errors including stop-codon readthrough, +1 frameshifting, and −1 frameshifting (49, 83). Mapping 2’-O-methylation changes arising from fibrillarin knockdown onto the ribosome structure revealed that altered 2’-O-methylations can be found in several regions involved in intermolecular interactions, such as between tRNA and the A-site, inter-subunit bridges, or around the peptide exit tunnel (27). While these changes in 2’-O-methylation levels did not affect translation elongation, they affected the IRES-mediated translation initiation. Our data reveal that changes in 2’-O-methylation levels can affect translation fidelity in multiple ways. In addition to modulating IRES-dependent translation initiation, similar to what was observed after fibrillarin knockdown, we also observed changes in the rate of frameshifting and near-cognate start codon recognition. Frameshifting involves the pause of the ribosomal subunits in the rotated state prior to translocation (59–61). Our data indicate that hypo 2’-O-methylated ribosomes from *bcd1-D72A* cells assume more rotated state *in vivo* (Figure 3A-F). Because we observed a similar preference in rotational state of ribosomes in Nop1 deficient cells (Figure 3G, H), our data suggest that the observed changes in rRNA dynamics are mainly due to the changes in rRNA 2’-O-methylation pattern and not the rRNA folding defects resulting from the reduced snoRNA levels.

Previously, global pseudouridylation defects were shown to affect the binding of the A- and P-site tRNAs to the ribosome, explaining the increased frameshifting and decreased stop-codon readthrough rates of such ribosomes (11). Mapping of the hypomethylation sites around the E- and P-site tRNAs did not reveal dramatic changes in the methylation pattern, with the exception of two sites near the acceptor arm of each tRNA (Figure S4 A, B). We also did not observe a change in the rate of HPG incorporation into newly synthesized proteins, despite its overall lower incorporation at any time point (Figure 2A), suggesting that the translation rate remains unchanged. These findings corroborate the previous observation that hypo 2’-O-methylation does not affect the elongation rate (27). Whether long-range effects from hypo 2’-O-methylated sites can also influence tRNA binding remains to be addressed.

To our knowledge, the effect of rRNA 2’-O-methylations on the near-cognate start codon selection was unknown to date. Here, we show that eIF1, a major antagonist of near-cognate start site selection, has a lower affinity for hypomethylated 40S than wild type 40S (Figure 4A-C). Interestingly, overexpression of eIF1, but not eIF1A, suppressed the slow growth phenotype of *bcd1-D72A* cells to a great extent without affecting the growth of wild-type control (Figure 4D). A point mutation in the *RPS3* gene which hampers the near-cognate start codon selection also rescued the slow growth phenotype of *bcd1-D72A*, albeit to a lesser extent than eIF1 (Figure 4E). Collectively, these results suggest that a major role of 2’-O-methylation of ribosomes is to support faithful translation initiation. The increased rate of near-cognate start site selection results in the production of peptides from many short open reading frames (84) or the production of proteins with extended N-termini (85). Recently, near-cognate start site selection was also linked to the re-distribution of many proteins from the cytosol to mitochondria due to the gain of N-terminal mitochondrial targeting signals (86). It is not yet clear whether rRNA hypo 2’-O-methylation causes similar defects. As we did not detect major changes in the 2’-O-methylation of rRNA in the vicinity of the eIF1 binding site (Figure S4C), we speculate that the decreased affinity of eIF1 for the ribosome is due to a change in the dynamics of the 40S ribosomal subunit (87). According to this model, in vacant 40S the head and body fluctuate relative to each other, and eIF1 preferentially binds and stabilizes the open conformation. In *bcd1-D72A* cells, the equilibrium between open and closed conformations is altered (Figure 4F, G), resulting in weaker binding of eIF1 to the 40S subunit. Future studies are needed to test this hypothesis. Taken together, our data reveal an unforeseen spectrum of 2’-O-methylation heterogeneity in yeast rRNA and suggest a significant role for rRNA 2’-O-methylation in regulating cellular translation by controlling ribosome dynamics and ligand binding.

## Materials and Methods

### Plasmids and strains

Plasmids used in this study are listed in Table S3. All mutations were introduced by site-directed mutagenesis and confirmed by sequencing. Strains are listed in Table S4. All yeast strains were confirmed by PCR followed by sequencing, as well as western blotting when antibodies were available.

### RiboMeth-seq

RiboMeth-Seq was essentially performed as previously reported (31). Briefly, 150 ng total RNA was fragmented under denaturing conditions using an alkaline buffer (pH 9.4) to obtain average size of 20-40 nt. Fragments were end-repaired and ligated to adapters using NEBNext Small RNA kit for Illumina. Sequencing was performed on Illumina HiSeq1000. Reads were mapped to the yeast rDNA and snoRNA sequences, and the RiboMeth-Seq score (fraction methylated) was calculated as MethScore (for +/− 2 nt) (88), equivalent to “ScoreC” (30). Statistical significance was determined using Student’ s t-test (P < 0.05).

### HPG incorporation assay

BY4741 and *bcd1-D72A* cells were transformed with pRS411 and grown in synthetic media lacking methionine at 30°C to mid-log phase. L-Homopropargylglycine (HPG) was added to 10 mL cultures to a final concentration of 50 μM and cells were incubated at 30°C. At each indicated time point, 2 ml of the culture was removed, and cells were washed with cold water and frozen in liquid nitrogen. The cell pellets were resuspended in 100 μL of RIPA buffer (50mM Tris-HCl, pH 7.4, 150mM sodium chloride, 1mM EDTA, 1mM EGTA, 1% Triton X-100, 1% sodium deoxycholate and 0.1% SDS) supplemented with protease inhibitors, mixed with disruption beads, and lysed in a bead beater. After clearing the lysate, the protein concentration was measured using BCA assay (ThermoFisher), and an equal amount of protein was used for labeling by Alexa Fluor 488 using the Click-iT™ HPG Alexa Fluor™ 488 Protein Synthesis Assay Kit (ThermoFisher). Labeled proteins were resolved on 12% SDS gel and visualized on a ChemiDoc Imager (BioRad). Total protein was visualized after Coomassie staining and imaged using ChemiDoc.

### Fidelity assay

Translation fidelity was measured using previously established dual-luciferase reporters (54–58). For measurements, 1 mL of BY4741 and *bcd1-D72A* cells expressing the dual luciferase plasmids were pelleted in the mid-log phase, washed, and frozen in liquid nitrogen. Luciferase activities were measured using the Promega Dual-Luciferase kit by resuspending cells in 200mL of 1X Passive Lysis Buffer and incubating for 5 min. To measure firefly and Renilla activities, 10 μL lysates were mixed with 30 μL Luciferase Assay Reagent II in white, clear bottom 96W Microplates (Costar), followed by an addition of 30 μL Stop&Glo Reagent. Measurements were performed using a Synergy Microplate reader (BioTek). For each sample, firefly luciferase activity was normalized against Renilla activity, and then values observed for *bcd1-D72A* were normalized against wild-type control.

### *In vivo* RNA structure probing

BY4741 and *bcd1-D72A* cells were grown to the mid-log phase at 30°C. SSU-G913 was probed using phenylglyoxal. The cultures were divided into two tubes, mixed with either phenylglyoxal (16 mM final concentration) or an equal volume of DMSO, and incubated for 5 min at 30°C before washing with water. RNA was extracted using phenol/chloroform and precipitated with ethanol. Precipitated RNA was resuspended in water and treated with DNase I (BioRad) and further purified with the Quick RNA miniprep kit (Zymo Research) before reverse transcription was performed using Superscript III (ThermoFisher) according to the manufacturer’ s protocol. To probe SSU-G913 in *nop1-ts* cells, experimental ts cells and wild-type control cells were grown to mid-log phase at 37°C before treatment with phenylglyoxal as before. Data were quantified using Image Lab Software (Biorad). The intensity of bands at the reverse transcription stops were normalized to all band intensities in each lane.

### *In vitro* RNA structure probing

25 nM purified 40S ribosomes from control or *bcd1-D72A* cells was mixed with 200 mM DMS or ethanol in 80 mM Hepes/NaOH (pH 7.4), 50 mM NaCl and 0.5 mM MgOAc. The mixture was incubated at 30°C for 5 minutes before quenching with 400 mM betamercaptoethanol/600 mM NaOAc. The RNA was precipitated with ethanol and further purified using a Quick RNA miniprep kit (Zymo Research) before reverse transcription was performed using Superscript III (ThermoFisher) according to the manufacturer’ s protocol. Data were quantified using Image Lab Software (Biorad). To quantify nucleotide accessibility and account for loading differences, the intensity of bands at the reverse transcription stops were normalized to all band intensities in each lane.

### Purification of 40S and eIF1

40S ribosomes from BY4741 and BCD1-D72A cells were purified as previously described (89). eIF1 was expressed as a 6xHis-tagged protein in Rosetta2(DE3) cells from a pET23-6xHis-TEV-eIF1 plasmid. Protein expression was induced at OD_600_ ~0.6 by addition of 1 mM IPTG and the cultures were incubated for 16 hours at 18°C. eIF1 was purified on Ni-NTA resin in buffer A (500 mM NaCl, 50 mM HEPES-NaOH pH 7.5, 20 mM imidazole and 5% glycerol). The protein was eluted with 240 mM imidazole and further purified over a Superdex S-75 gel filtration column (GE Healthcare) equilibrated in 150 mM NaCl, 20 mM HEPES-NaOH pH 7.5, 5 % glycerol, and 1 mM DTT.

### eIF1-40S binding assay

6xHis-eIF1 was labeled with a Cy5 fluorophore through non-covalent linkage to the His-tag moiety using the Monolith Protein Labeling Kit RED-tris-NTA 2^nd^ generation (NanoTemper Technologies). 40S subunits (0.58 nM –1.2 μM) isolated from wild-type control or *bcd1-D72A* cells were incubated with 100 nM labeled eIF1. Fluorescence was measured using a Dianthus NT.23 Pico instrument (NanoTemper Technologies). Fluorescence values were baseline corrected and plotted against 40S concentration. Data were fitted with a non-linear regression model in GraphPad Prism 8.0.

### Growth assay

Cells grown to mid-log phase in minimal media were diluted into fresh media and growth rates were measured in the Epoch2 microplate reader (BioTek) by recording OD_600_ every 20 min.

### Analysis of the steady-state levels of snoRNAs

*NOP1* or *nop1-ts* cells were grown in YPD at 37°C to OD_600_ ~ 0.6. Total RNA from three biological replicates of each strain was isolated using the hot phenol method. snoRNAs were separated on 8% acrylamide/urea gels, transferred to Hybond nylon membrane (GE Healthcare), and probed as indicated. Bands were quantified using Image Lab software (Bio-Rad).

## Acknowledgments

This work was supported by startup funds from Emory University and NIH Grant 1R35GM138123-01 (to H.G.). R. E. D-W. was supported by an NSF Graduate Research Fellowship. We thank Drs. David Bedwell, John Dinmann, Alan Hinnebusch, Sunnie Thompson for the gift of plasmids and strains. We also thank members of the Ghalei lab, and Drs. Anita Corbett and Daniel Reines for comments on the manuscript.

## References

1. M. Helm, Post-transcriptional nucleotide modification and alternative folding of RNA. Nucleic acids research 34, 721–733 (2006).

2. I. A. Roundtree, M. E. Evans, T. Pan, C. He, Dynamic RNA Modifications in Gene Expression Regulation. Cell 169, 1187–1200 (2017).

3. L. Ayadi, A. Galvanin, F. Pichot, V. Marchand, Y. Motorin, RNA ribose methylation (2’-O-methylation): Occurrence, biosynthesis and biological functions. Biochim Biophys Acta Gene Regul Mech 1862, 253–269 (2019).

4. K. E. Sloan et al., Tuning the ribosome: The influence of rRNA modification on eukaryotic ribosome biogenesis and function. RNA Biol 14, 1138–1152 (2017).

5. P. L. Monaco, V. Marcel, J. J. Diaz, F. Catez, 2’-O-Methylation of Ribosomal RNA: Towards an Epitranscriptomic Control of Translation? Biomolecules 8 (2018).

6. S. K. Natchiar, A. G. Myasnikov, H. Kratzat, I. Hazemann, B. P. Klaholz, Visualization of chemical modifications in the human 80S ribosome structure. Nature 551, 472–477 (2017).

7. W. A. Decatur, M. J. Fournier, rRNA modifications and ribosome function. Trends Biochem Sci 27, 344–351 (2002).

8. D. G. Dimitrova, L. Teysset, C. Carre, RNA 2’-O-Methylation (Nm) Modification in Human Diseases. Genes (Basel) 10 (2019).

9. S. K. Natchiar, A. G. Myasnikov, I. Hazemann, B. P. Klaholz, Visualizing the Role of 2’-OH rRNA Methylations in the Human Ribosome Structure. Biomolecules 8 (2018).

10. Y. S. Polikanov, S. V. Melnikov, D. Söll, T. A. Steitz, Structural insights into the role of rRNA modifications in protein synthesis and ribosome assembly. Nat Struct Mol Biol 22, 342–344 (2015).

11. K. Jack et al., rRNA pseudouridylation defects affect ribosomal ligand binding and translational fidelity from yeast to human cells. Mol Cell 44, 660–666 (2011).

12. J. Jiang, H. Seo, C. S. Chow, Post-transcriptional Modifications Modulate rRNA Structure and Ligand Interactions. Acc Chem Res 49, 893–901 (2016).

13. H. Abou Assi et al., 2’-O-Methylation can increase the abundance and lifetime of alternative RNA conformational states. Nucleic Acids Res 48, 12365–12379 (2020).

14. P. Auffinger, E. Westhof, Rules governing the orientation of the 2’-hydroxyl group in RNA. J Mol Biol 274, 54–63 (1997).

15. G. Kawai et al., Conformational rigidity of specific pyrimidine residues in tRNA arises from posttranscriptional modifications that enhance steric interaction between the base and the 2’-hydroxyl group. Biochemistry 31, 1040–1046 (1992).

16. E. Malek-Adamian et al., Adjusting the Structure of 2’-Modified Nucleosides and Oligonucleotides via C4’-α-F or C4’-α-OMe Substitution: Synthesis and Conformational Analysis. J Org Chem 83, 9839–9849 (2018).

17. D. M. Cheng, R. H. Sarma, Nuclear magnetic resonance study of the impact of ribose 2’-O-methylation on the aqueous solution conformation of cytidylyl-(3’ leads to 5’)-cytidine. Biopolymers 16, 1687–1711 (1977).

18. S. K. Mahto, C. S. Chow, Synthesis and solution conformation studies of the modified nucleoside N(4),2’-O-dimethylcytidine (m(4)Cm) and its analogues. Bioorg Med Chem 16, 8795–8800 (2008).

19. C. S. Chow, T. N. Lamichhane, S. K. Mahto, Expanding the nucleotide repertoire of the ribosome with post-transcriptional modifications. ACS Chem Biol 2, 610–619 (2007).

20. Z. Kiss-Laszlo, Y. Henry, J. P. Bachellerie, M. Caizergues-Ferrer, T. Kiss, Site-specific ribose methylation of preribosomal RNA: a novel function for small nucleolar RNAs. Cell 85, 1077–1088 (1996).

21. J. P. Bachellerie, J. Cavaillé, Guiding ribose methylation of rRNA. Trends Biochem Sci 22, 257–261 (1997).

22. G. Yu, Y. Zhao, H. Li, The multistructural forms of box C/D ribonucleoprotein particles. Rna 24, 1625–1633 (2018).

23. A. Lapinaite et al., The structure of the box C/D enzyme reveals regulation of RNA methylation. Nature 502, 519–523 (2013).

24. J. Lin et al., Structural basis for site-specific ribose methylation by box C/D RNA protein complexes. Nature 469, 559–563 (2011).

25. N. J. Watkins, M. T. Bohnsack, The box C/D and H/ACA snoRNPs: key players in the modification, processing and the dynamic folding of ribosomal RNA. Wiley interdisciplinary reviews. RNA 3, 397–414 (2012).

26. S. Massenet, E. Bertrand, C. Verheggen, Assembly and trafficking of box C/D and H/ACA snoRNPs. RNA Biol 14, 680–692 (2017).

27. J. Erales et al., Evidence for rRNA 2’-O-methylation plasticity: Control of intrinsic translational capabilities of human ribosomes. Proc Natl Acad Sci U S A 114, 12934–12939 (2017).

28. S. Sharma, V. Marchand, Y. Motorin, D. L. J. Lafontaine, Identification of sites of 2’-O-methylation vulnerability in human ribosomal RNAs by systematic mapping. Sci Rep 7, 11490 (2017).

29. J. Yang et al., Mapping of Complete Set of Ribose and Base Modifications of Yeast rRNA by RP-HPLC and Mung Bean Nuclease Assay. PloS one 11, e0168873 (2016).

30. U. Birkedal et al., Profiling of ribose methylations in RNA by high-throughput sequencing. Angew Chem Int Ed Engl 54, 451–455 (2015).

31. V. Marchand, F. Blanloeil-Oillo, M. Helm, Y. Motorin, Illumina-based RiboMethSeq approach for mapping of 2’-O-Me residues in RNA. Nucleic acids research 44, e135 (2016).

32. M. Taoka et al., Landscape of the complete RNA chemical modifications in the human 80S ribosome. Nucleic Acids Res 46, 9289–9298 (2018).

33. M. Taoka et al., The complete chemical structure of Saccharomyces cerevisiae rRNA: partial pseudouridylation of U2345 in 25S rRNA by snoRNA snR9. Nucleic acids research 44, 8951–8961 (2016).

34. N. Krogh et al., Profiling of 2’-O-Me in human rRNA reveals a subset of fractionally modified positions and provides evidence for ribosome heterogeneity. Nucleic Acids Res 44, 7884–7895 (2016).

35. Y. Motorin, M. Quinternet, W. Rhalloussi, V. Marchand, Constitutive and variable 2’-O-methylation (Nm) in human ribosomal RNA. RNA Biol 10.1080/15476286.2021.1974750, 1–10 (2021).

36. M. B. Ferretti, K. Karbstein, Does functional specialization of ribosomes really exist? Rna 25, 521–538 (2019).

37. V. Marcel et al., Ribosomal RNA 2’O-methylation as a novel layer of inter-tumour heterogeneity in breast cancer. NAR Cancer 2, zcaa036 (2020).

38. J. Hebras, N. Krogh, V. Marty, H. Nielsen, J. Cavaillé, Developmental changes of rRNA ribose methylations in the mouse. RNA Biol 17, 150–164 (2020).

39. V. Marcel et al., p53 acts as a safeguard of translational control by regulating fibrillarin and rRNA methylation in cancer. Cancer Cell 24, 318–330 (2013).

40. Y. Zhao, J. Rai, H. Yu, H. Li, Pseudouridine-free Ribosome Exhibits Distinct Inter-subunit Movements. bioRxiv 10.1101/2021.06.02.446812, 2021.2006.2002.446812 (2021).

41. D. Tollervey, H. Lehtonen, M. Carmo-Fonseca, E. C. Hurt, The small nucleolar RNP protein NOP1 (fibrillarin) is required for pre-rRNA processing in yeast. The EMBO journal 10, 573–583 (1991).

42. D. Tollervey, H. Lehtonen, R. Jansen, H. Kern, E. C. Hurt, Temperature-sensitive mutations demonstrate roles for yeast fibrillarin in pre-rRNA processing, pre-rRNA methylation, and ribosome assembly. Cell 72, 443–457 (1993).

43. M. A. Amin et al., Fibrillarin, a nucleolar protein, is required for normal nuclear morphology and cellular growth in HeLa cells. Biochem Biophys Res Commun 360, 320–326 (2007).

44. F. J. LaRiviere, S. E. Cole, D. J. Ferullo, M. J. Moore, A late-acting quality control process for mature eukaryotic rRNAs. Mol Cell 24, 619–626 (2006).

45. P. Tessarz et al., Glutamine methylation in histone H2A is an RNA-polymerase-I-dedicated modification. Nature 505, 564–568 (2014).

46. W. T. Peng et al., A panoramic view of yeast noncoding RNA processing. Cell 113, 919–933 (2003).

47. A. Paul et al., Bcd1p controls RNA loading of the core protein Nop58 during C/D box snoRNP biogenesis. RNA (New York, N.Y.) 25, 496–506 (2019).

48. S. Khoshnevis, R. E. Dreggors, T. F. R. Hoffmann, H. Ghalei, A conserved Bcd1 interaction essential for box C/D snoRNP biogenesis. J Biol Chem 294, 18360–18371 (2019).

49. A. Baudin-Baillieu et al., Nucleotide modifications in three functionally important regions of the Saccharomyces cerevisiae ribosome affect translation accuracy. Nucleic acids research 37, 7665–7677 (2009).

50. X. H. Liang, Q. Liu, M. J. Fournier, rRNA modifications in an intersubunit bridge of the ribosome strongly affect both ribosome biogenesis and activity. Mol Cell 28, 965–977 (2007).

51. E. W. Mills, R. Green, Ribosomopathies: There’s strength in numbers. Science 358 (2017).

52. S. O. Sulima, I. J. F. Hofman, K. De Keersmaecker, J. D. Dinman, How Ribosomes Translate Cancer. Cancer Discov 7, 1069–1087 (2017).

53. Y. Yi et al., A PRC2-independent function for EZH2 in regulating rRNA 2’-O methylation and IRES-dependent translation. Nat Cell Biol 23, 341–354 (2021).

54. J. W. Harger, J. D. Dinman, An in vivo dual-luciferase assay system for studying translational recoding in the yeast Saccharomyces cerevisiae. RNA 9, 1019–1024 (2003).

55. Y. N. Cheung et al., Dissociation of eIF1 from the 40S ribosomal subunit is a key step in start codon selection in vivo. Genes Dev 21, 1217–1230 (2007).

56. K. M. Keeling et al., Leaky termination at premature stop codons antagonizes nonsense-mediated mRNA decay in S. cerevisiae. RNA 10, 691–703 (2004).

57. J. Salas-Marco, D. M. Bedwell, Discrimination between defects in elongation fidelity and termination efficiency provides mechanistic insights into translational readthrough. J Mol Biol 348, 801–815 (2005).

58. D. M. Landry, M. I. Hertz, S. R. Thompson, RPS25 is essential for translation initiation by the Dicistroviridae and hepatitis C viral IRESs. Genes Dev 23, 2753–2764 (2009).

59. P. Qin, D. Yu, X. Zuo, P. V. Cornish, Structured mRNA induces the ribosome into a hyper-rotated state. EMBO Rep 15, 185–190 (2014).

60. J. Choi, S. O’ Loughlin, J. F. Atkins, J. D. Puglisi, The energy landscape of −1 ribosomal frameshifting. Sci Adv 6, eaax6969 (2020).

61. J. Chen et al., Dynamic pathways of −1 translational frameshifting. Nature 512, 328–332 (2014).

62. S. O. Sulima et al., Eukaryotic rpL10 drives ribosomal rotation. Nucleic Acids Res 42, 2049–2063 (2014).

63. A. Meskauskas, J. D. Dinman, Ribosomal protein L3: gatekeeper to the A site. Mol Cell 25, 877–888 (2007).

64. R. Belardinelli, H. Sharma, F. Peske, W. Wintermeyer, M. V. Rodnina, Translocation as continuous movement through the ribosome. RNA Biol 13, 1197–1203 (2016).

65. M. C. Justice et al., Elongation factor 2 as a novel target for selective inhibition of fungal protein synthesis. J Biol Chem 273, 3148–3151 (1998).

66. H. Huang, K. Karbstein, Assembly factors chaperone ribosomal RNA folding by isolating helical junctions that are prone to misfolding. Proc Natl Acad Sci U S A 118 (2021).

67. M. Kofoed et al., An Updated Collection of Sequence Barcoded Temperature-Sensitive Alleles of Yeast Essential Genes. G3 (Bethesda) 5, 1879–1887 (2015).

68. D. Maag, C. A. Fekete, Z. Gryczynski, J. R. Lorsch, A conformational change in the eukaryotic translation preinitiation complex and release of eIF1 signal recognition of the start codon. Mol Cell 17, 265–275 (2005).

69. L. A. Passmore et al., The eukaryotic translation initiation factors eIF1 and eIF1A induce an open conformation of the 40S ribosome. Mol Cell 26, 41–50 (2007).

70. T. V. Pestova, V. G. Kolupaeva, The roles of individual eukaryotic translation initiation factors in ribosomal scanning and initiation codon selection. Genes & development 16, 2906–2922 (2002).

71. J. S. Nanda et al., eIF1 controls multiple steps in start codon recognition during eukaryotic translation initiation. J Mol Biol 394, 268–285 (2009).

72. D. Maag, J. R. Lorsch, Communication between eukaryotic translation initiation factors 1 and 1A on the yeast small ribosomal subunit. J Mol Biol 330, 917–924 (2003).

73. J. Dong et al., Rps3/uS3 promotes mRNA binding at the 40S ribosome entry channel and stabilizes preinitiation complexes at start codons. Proc Natl Acad Sci U S A 114, E2126–E2135 (2017).

74. A. G. Hinnebusch, The scanning mechanism of eukaryotic translation initiation. Annu Rev Biochem 83, 779–812 (2014).

75. A. G. Hinnebusch, Structural Insights into the Mechanism of Scanning and Start Codon Recognition in Eukaryotic Translation Initiation. Trends in biochemical sciences 42, 589–611 (2017).

76. J. L. Llácer et al., Conformational Differences between Open and Closed States of the Eukaryotic Translation Initiation Complex. Mol Cell 59, 399–412 (2015).

77. D. Nachmani et al., Germline NPM1 mutations lead to altered rRNA 2’-O-methylation and cause dyskeratosis congenita. Nat Genet 51, 1518–1529 (2019).

78. F. Zhou et al., AML1-ETO requires enhanced C/D box snoRNA/RNP formation to induce self-renewal and leukaemia. Nat Cell Biol 19, 844–855 (2017).

79. S. Ojha, S. Malla, S. M. Lyons, snoRNPs: Functions in Ribosome Biogenesis. Biomolecules 10 (2020).

80. S. Parker et al., Large-scale profiling of noncoding RNA function in yeast. PLoS Genet 14, e1007253 (2018).

81. L. N. Balarezo-Cisneros et al., Functional and transcriptional profiling of non-coding RNAs in yeast reveal context-dependent phenotypes and in trans effects on the protein regulatory network. PLoS Genet 17, e1008761 (2021).

82. J. Esguerra, J. Warringer, A. Blomberg, Functional importance of individual rRNA 2’-O-ribose methylations revealed by high-resolution phenotyping. Rna 14, 649–656 (2008).

83. X. H. Liang, Q. Liu, M. J. Fournier, Loss of rRNA modifications in the decoding center of the ribosome impairs translation and strongly delays pre-rRNA processing. RNA 15, 1716–1728 (2009).

84. S. D. Kulkarni et al., Temperature-dependent regulation of upstream open reading frame translation in S. cerevisiae. BMC Biol 17, 101 (2019).

85. F. Zhou, H. Zhang, S. D. Kulkarni, J. R. Lorsch, A. G. Hinnebusch, eIF1 discriminates against suboptimal initiation sites to prevent excessive uORF translation genome-wide. RNA 26, 419–438 (2020).

86. G. Monteuuis et al., Non-canonical translation initiation in yeast generates a cryptic pool of mitochondrial proteins. Nucleic Acids Res 47, 5777–5791 (2019).

87. D. Maag, M. A. Algire, J. R. Lorsch, Communication between eukaryotic translation initiation factors 5 and 1A within the ribosomal pre-initiation complex plays a role in start site selection. Journal of molecular biology 356, 724–737 (2006).

88. F. Pichot et al., Holistic Optimization of Bioinformatic Analysis Pipeline for Detection and Quantification of 2’-O-Methylations in RNA by RiboMethSeq. Front Genet 11, 38 (2020).

89. M. G. Acker, S. E. Kolitz, S. F. Mitchell, J. S. Nanda, J. R. Lorsch, Reconstitution of yeast translation initiation. Methods Enzymol 430, 111–145 (2007).

