## Supplemental Figures and Tables for "Ribosomal RNA 2’-O-methylations regulate translation by impacting ribosome dynamics"

**Supplementary Figures and Tables**

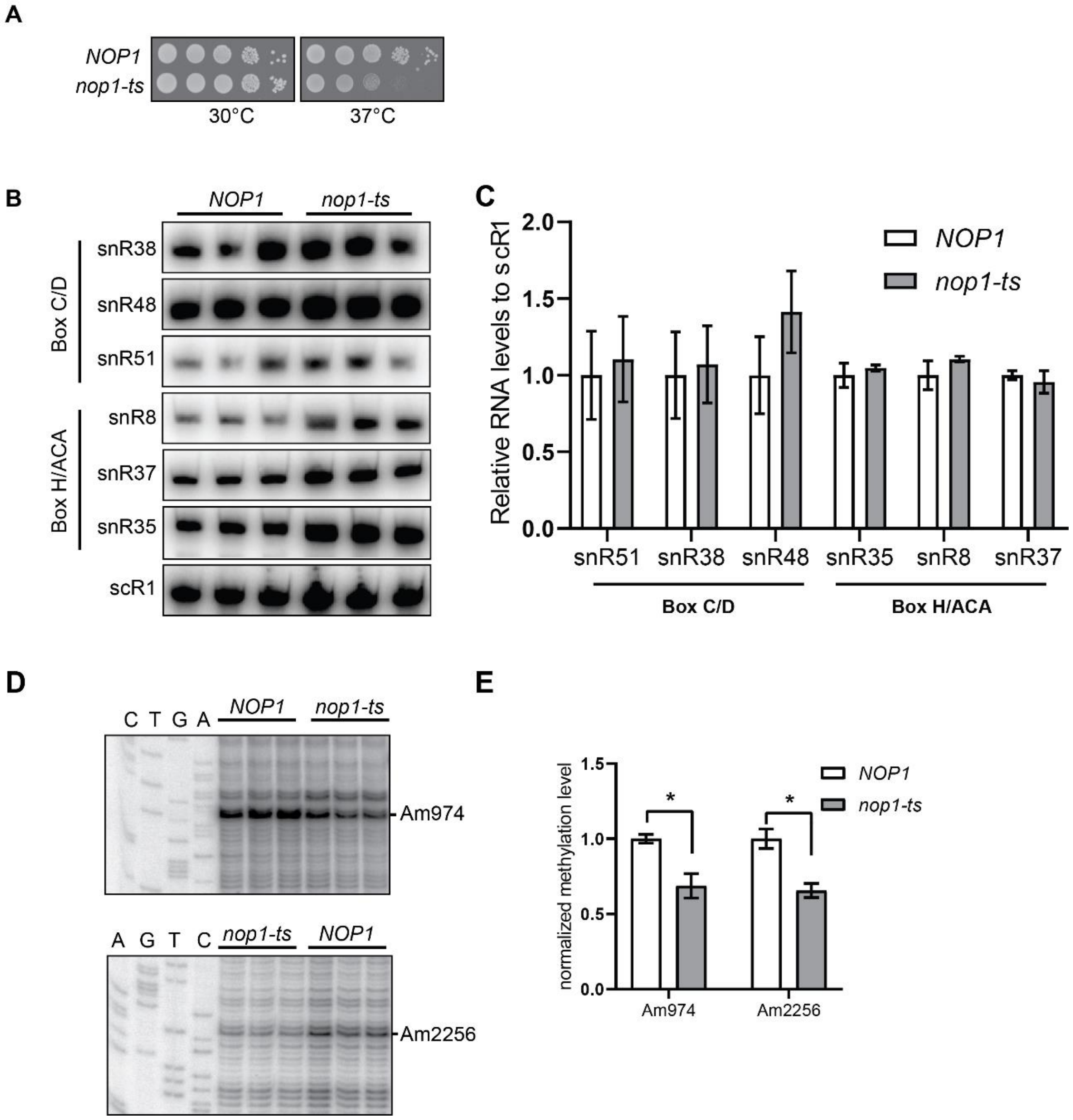


**Figure S1. *nop1-ts* affects the rRNA 2'-O methylation but not the snoRNA levels. (A)** *NOP1* and *nop1-ts* cells were serially diluted and spotted on YPD plates and incubated at 30°C or 37°C for two days. **(B)** Northern blotting analyses of steady-state expression levels of box C/D and box H/ACA snoRNAs in *NOP1* and *nop1-ts* cells. **(C)** Quantification of data shown in A as normalized to scR1. Graph bars represent the mean and S.D. from three biological replicates. There was no significant difference in the snoRNA levels between *NOP1* and *nop1-ts* cells as determined using an unpaired t test. **(D)** Reverse transcription at low concentration of dNTP combined with sequencing gel analysis was used to determine the methylation levels at a methylation site in 18S rRNA (Am947) and 25S rRNA (Am2256). **(E)** quantification of the data in D. Graph bars represent the mean and S.D. from three biological replicates.

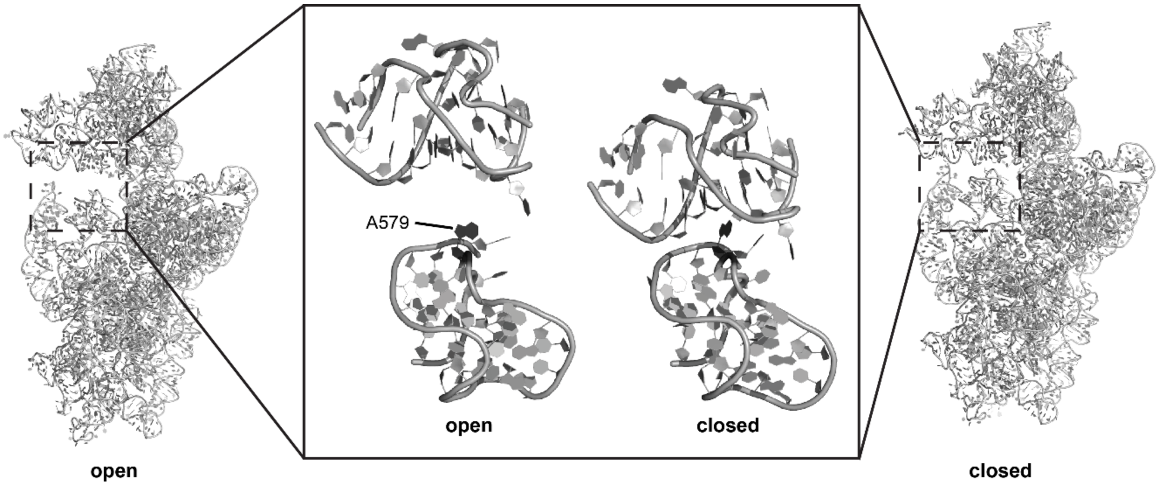


**Figure S2. Comparing the position of SSU-A579 in the open and closed conformations.** The left and right structures are the 18S rRNA in the open (PDB: 3jaq) and closed (PDB: 3jap) conformations, respectively. The position of the mRNA latch is boxed in both structures and zoomed in in the middle panel.

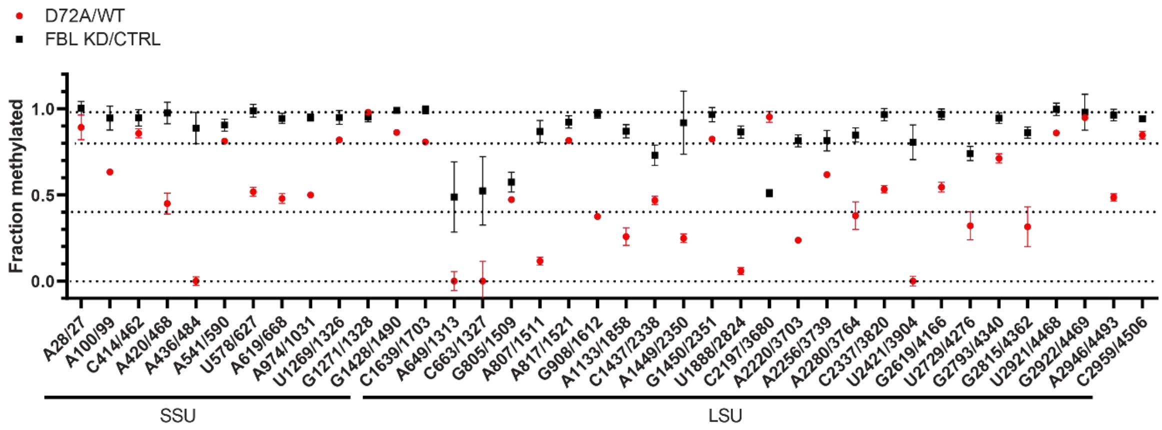


**Figure S3. Comparison of the changes in rRNA 2'-O-methylation levels of the sites conserved between yeast and human.** The red squares show mean methylation scores at each indicated site in bcd1-D72A cells relative to control wild-type yeast. The black squares show mean methylation scores from HEK cells after fibrillarin knockdown relative to control (1). Error bars represent SD.

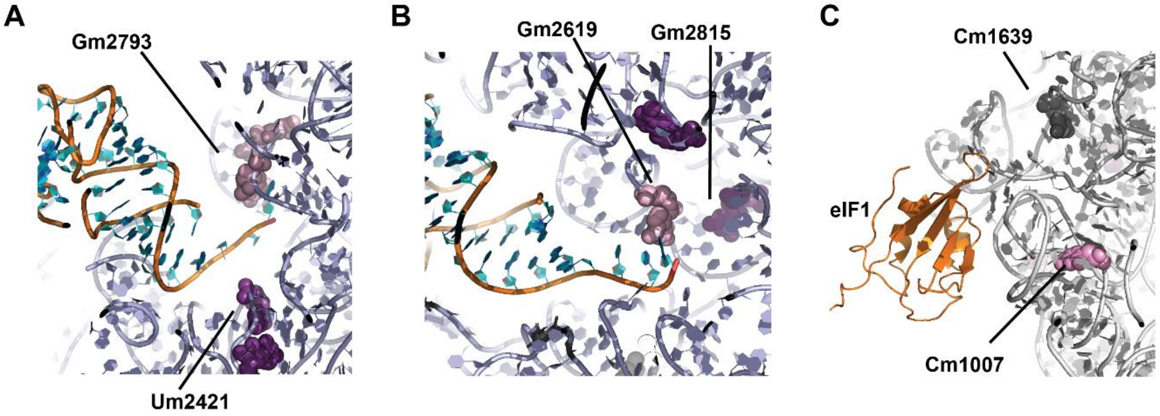


**Figure S4. Position of rRNA 2'-O-methylations relative to the E- and P-site tRNAs and eIF1.** (**A**) The E-site tRNA (**B**) the P-site tRNA (PDB: 3j78). The 25S rRNA is in light blue and the tRNA in orange. Methylation sites follow the same color coding as in Figure 1. (**C**) eIF1 binding site (PDB: 6gsm). The 18S rRNA is in grey and the tRNA in orange.

**Table S1: Quantitative mapping of rRNA 2′-O-Me in yeast cells.** The 2′-O-methylation levels at each site of 18S and 25S rRNA was evaluated by RiboMeth-Seq for wild-type control and mutant *bcd1-D72A* yeast cell. Data are expressed as mean MethScore (Score C) values ± SD (n=3 independent biological replicates) for each known methylated nucleotide in yeast rRNA.

| 18S | WT | SD | D72A | SD | 25S | WT | SD | D72A | SD |
|---|---|---|---|---|---|---|---|---|---|
| A28 | 0.89 | 0.003 | 0.80 | 0.030 | A649 | 0.83 | 0.008 | -0.05 | 0.022 |
| A100 | 0.81 | 0.003 | 0.51 | 0.007 | C650 | 0.87 | 0.009 | 0.67 | 0.007 |
| C414 | 0.85 | 0.010 | 0.73 | 0.010 | C663 | 0.48 | 0.018 | -0.53 | 0.026 |
| A420 | 0.80 | 0.014 | 0.36 | 0.023 | G805 | 0.87 | 0.001 | 0.41 | 0.005 |
| A436 | 0.65 | 0.027 | -0.02 | 0.008 | A807 | 0.81 | 0.006 | 0.09 | 0.009 |
| A541 | 0.89 | 0.003 | 0.73 | 0.005 | A817 | 0.91 | 0.005 | 0.74 | 0.006 |
| G562 | 0.81 | 0.001 | 0.62 | 0.008 | G867 | 0.83 | 0.003 | 0.49 | 0.012 |
| U578 | 0.84 | 0.005 | 0.43 | 0.010 | A876 | 0.86 | 0.004 | 0.61 | 0.009 |
| A619 | 0.87 | 0.005 | 0.42 | 0.012 | U898 | 0.76 | 0.004 | 0.16 | 0.008 |
| A796 | 0.88 | 0.003 | 0.17 | 0.016 | G908 | 0.87 | 0.006 | 0.33 | 0.007 |
| A974 | 0.89 | 0.003 | 0.45 | 0.007 | A1133 | 0.87 | 0.001 | 0.22 | 0.021 |
| C1007 | 0.88 | 0.004 | 0.41 | 0.011 | C1437 | 0.89 | 0.003 | 0.42 | 0.010 |
| G1126 | 0.82 | 0.002 | 0.60 | 0.003 | A1449 | 0.84 | 0.006 | 0.21 | 0.010 |
| U1269 | 0.89 | 0.006 | 0.73 | 0.002 | G1450 | 0.91 | 0.001 | 0.75 | 0.003 |
| G1271 | 0.85 | 0.004 | 0.83 | 0.004 | U1888 | 0.82 | 0.002 | 0.05 | 0.008 |
| G1428 | 0.87 | 0.002 | 0.75 | 0.002 | C2197 | 0.77 | 0.009 | 0.74 | 0.011 |
| G1572 | 0.85 | 0.013 | 0.07 | 0.011 | A2220 | 0.94 | 0.003 | 0.22 | 0.007 |
| C1639 | 0.89 | 0.003 | 0.72 | 0.004 | A2256 | 0.79 | 0.008 | 0.49 | 0.002 |
| | | | | | A2280 | 0.90 | 0.007 | 0.34 | 0.034 |
| | | | | | A2281 | 0.87 | 0.010 | 0.52 | 0.028 |
| | | | | | G2288 | 0.93 | 0.003 | 0.56 | 0.015 |
| | | | | | C2337 | 0.89 | 0.002 | 0.48 | 0.009 |
| | | | | | U2417 | 0.87 | 0.005 | 0.14 | 0.017 |
| | | | | | U2421 | 0.91 | 0.004 | -0.13 | 0.012 |
| | | | | | G2619 | 0.92 | 0.005 | 0.50 | 0.012 |
| | | | | | A2640 | 0.87 | 0.002 | 0.56 | 0.014 |
| | | | | | U2724 | 0.94 | 0.003 | 0.52 | 0.011 |
| | | | | | U2729 | 0.76 | 0.012 | 0.24 | 0.029 |
| | | | | | G2791 | 0.81 | 0.011 | 0.58 | 0.013 |
| | | | | | G2793 | 0.88 | 0.006 | 0.63 | 0.011 |
| | | | | | G2815 | 0.85 | 0.004 | 0.27 | 0.046 |
| | | | | | U2921 | 0.93 | 0.006 | 0.80 | 0.008 |
| | | | | | A2946 | 0.89 | 0.008 | 0.43 | 0.009 |
| | | | | | C2948 | 0.56 | 0.016 | 0.03 | 0.009 |
| | | | | | C2959 | 0.89 | 0.003 | 0.76 | 0.009 |

**Table S2: Lack of correlation between the snoRNA and 2'-O methylation levels.** Comparison of the snoRNA levels corresponding to the most stable 2'-O-methylation sites (mean C score > 0.8) in 18S and 25S rRNA.

| | **2'-O-methylation site** | | **Box C/D snoRNA level** | |
|---|---|---|---|---|
| **18S** | A28 | 0.89 | snR47 | 0.12 |
| | C414 | 0.86 | snR128 | 0.30 |
| | A541 | 0.81 | snR41 | 0.24 |
| | U1269 | 0.82 | snR65 | 0.20 |
| | G1271 | 0.98 | snR40 | 0.34 |
| | G1428 | 0.86 | snR56 | 0.13 |
| | C1639 | 0.81 | snR70 | 0.17 |
| **25S** | A817 | 0.82 | snR60 | 0.17 |
| | C1450 | 0.82 | snR24 | 0.32 |
| | C2197 | 0.95 | snR76 | 0.75 |
| | U2921 | 0.86 | snR52 | 0.09 |
| | C2959 | 0.85 | snR73 | 0.18 |

**Table S3**: **Plasmids used in this work**

| **Plasmid** | **Description** | **Backbone** | **Reference** |
|---|---|---|---|
| pDH412 | RPS3-WT | pRS315 | (2) |
| pDH425 | RPS3-R116D | pRS315 | (2) |
| pDH482 | PRS3-R117D | pRS315 | (2) |
| HG327 | RPL3-W255C | pRS425 | This study |
| HG3274 | RPL3-H256A | pRS425 | This study |
| HG3209 | SUI1+UTRs | pRS425 | This study |
| HG3210 | TIF11+UTRs | pRS425 | This study |
| pSRT209 | dicistronic reporter with a wild-type CrPV IGR IRES and a ΔAUG for the luciferase codon that reduces background from cryptic promoters | pRS425 | (3) |

| | | | |
|---|---|---|---|
| pSRT210 | identical to pSRT209 except for the 2 nucleotides change in the IRES to disrupt PKI | pRS425 | (3) |
| pDB722 | PGK-Readthrough | pYEplac195 | (4) |
| pDB723 | PGK- Stop | pYEplac195 | (4) |
| pDB868 | PGK- Miscoding Firefly H245R | pYEplac195 | (5) |
| pJD375 | ADH-0-Frame Control | pRS416 | (6) |
| pJD376 | ADH L-A (-1) Frameshift | pRS416 | (6) |
| pJD377 | ADH Ty1 (+1) Frameshift | pRS416 | (6) |
| pRaugFuug | RaugFuug ADH/GPD start | pRS416 | (7) |
| pRaugFaug | RaugFaug ADH/GPD start | pRS416 | (7) |
| HG187 | 6xHis-TEV-eIF1 | pET23a | This study |

**Table S4**: **Yeast strains used in this work**

| **Strain name** | **Description** | **Background** | **Genotype** | **Reference** |
|---|---|---|---|---|
| yHG000 | Wild-type | BY4741 | *MATα his3Δ1 leu2Δ0 met15Δ0 ura3Δ0* | GE Dharmacon |
| yHG458 | *bcd1-D72A* | BY4741 | *MATα his3Δ1 leu2Δ0 met15Δ0 ura3Δ0* | (8) |
| yHG520 | *ΔRPL3+rpl3-W255C* | BY4741 | *MATα his3Δ1 leu2Δ0 met15Δ0 ura3Δ0 RPL3::NAT* | This work |
| yHG521 | *ΔRPL3+rpl3-H256A* | BY4741 | *MATα his3Δ1 leu2Δ0 met15Δ0 ura3Δ0 RPL3::NAT* | This work |
| yHG561 | *Gal:RPS3* | BY4741 | *MATα his3Δ1 leu2Δ0 met15Δ0 ura3Δ0 GALRPS3::KAN* | This work |
| yHG562 | *bcd1-D72A* *Gal:RPS3* | BY4741 | *MATα his3Δ1 leu2Δ0 met15Δ0 ura3Δ0 GALRPS3::KAN* | This work |

**Supplementary References**


1. J. Erales *et al.*, Evidence for rRNA 2'-O-methylation plasticity: Control of intrinsic translational capabilities of human ribosomes. *Proc Natl Acad Sci U S A* **114**, 12934-12939 (2017).
2. J. Dong *et al.*, Rps3/uS3 promotes mRNA binding at the 40S ribosome entry channel and stabilizes preinitiation complexes at start codons. *Proc Natl Acad Sci U S A* **114**, E2126-E2135 (2017).
3. D. M. Landry, M. I. Hertz, S. R. Thompson, RPS25 is essential for translation initiation by the Dicistroviridae and hepatitis C viral IRESs. *Genes Dev* **23**, 2753-2764 (2009).
4. K. M. Keeling *et al.*, Leaky termination at premature stop codons antagonizes nonsense-mediated mRNA decay in S. cerevisiae. *RNA* **10**, 691-703 (2004).
5. J. Salas-Marco, D. M. Bedwell, Discrimination between defects in elongation fidelity and termination efficiency provides mechanistic insights into translational readthrough. *J Mol Biol* **348**, 801-815 (2005).
6. J. W. Harger, J. D. Dinman, An in vivo dual-luciferase assay system for studying translational recoding in the yeast Saccharomyces cerevisiae. *RNA* **9**, 1019-1024 (2003).
7. Y. N. Cheung *et al.*, Dissociation of eIF1 from the 40S ribosomal subunit is a key step in start codon selection in vivo. *Genes Dev* **21**, 1217-1230 (2007).
8. S. Khoshnevis, R. E. Dreggors, T. F. R. Hoffmann, H. Ghalei, A conserved Bcd1 interaction essential for box C/D snoRNP biogenesis. *J Biol Chem* **294**, 18360-18371 (2019).
